# Preclinical trial supports dual inhibition of BCL2 and Aurora kinase A for *MYCN*-amplified high-risk neuroblastoma

**DOI:** 10.64898/2026.09.06.749004

**Authors:** Alvin Kamili, Brandon A. Hearn, Isabella Temelkoska, Joan Solomon, Madeleine S. Wheatley, Christina H.T. Bui, Caroline J. Atkinson, Leith-Justin Elhassadi, Sadia Qureshi, Jayne Murray, Roxanne Cadiz, Andrew J. Gifford, Louise Cui, Lewis Crowley, Angela Lin, Chelsea Mayoh, Marie Wong-Erasmus, Timothy W. Failes, Greg M. Arndt, Michelle Haber, Murray D. Norris, M. Emmy M. Dolman, Toby N. Trahair, Jamie I. Fletcher

## Abstract

**Purpose:** Treatment for children with high-risk neuroblastoma relies on conventional chemotherapy and anti-GD2 immunotherapy. However, 5-year survival is only 50%, with high rates of late effects. Targeted therapy combinations are a major priority for these patients. The BCL2 inhibitor venetoclax, in combination with cyclophosphamide/topotecan, has clinical activity in relapsed and refractory neuroblastoma. We sought more effective and safer venetoclax combinations through systematic preclinical testing.

**Experimental design:** Synergistic combinations were identified by high-throughput screening using patient-derived xenograft (PDX) models and confirmed *in vivo*. The leading combination (venetoclax-alisertib) was compared to combination chemotherapy in a clinical trial-like study using 22 PDX models, in scheduling experiments designed to reduce short-term toxicity, and in combination with anti-GD2 immunotherapy. BCL2 and Bim-BCL2 complex protein levels were assessed as predictors of sensitivity.

**Results:** *In vitro* synergy with venetoclax was observed for standard-of-care chemotherapies and targeted agents, including DNA topoisomerase, microtubule, HDAC and Aurora kinase A (AURKA) inhibitors. Venetoclax-alisertib was particularly effective *in vivo*. In an n=1 study, venetoclax-alisertib induced objective response in all models. Activity was most striking in models of *MYCN*-amplified disease (n=12), doubling median survival time compared to cyclophosphamide/topotecan, and outperforming venetoclax-cyclophosphamide-topotecan. Efficacy was maintained with discontinuous schedules, minimizing hematological toxicity without substantially compromising activity. PDX-engrafted animals treated with venetoclax-alisertib and anti-GD2 immunotherapy survived tumor-free long-term. BCL2 expression and BCL2-Bim complex levels were of limited value for predicting response.

**Conclusion:** Our findings support advancement of BCL2-AURKA inhibition to clinical trial for neuroblastoma with or without anti-GD2 immunotherapy, particularly in patients with *MYCN* amplified disease.

**TRANSLATIONAL RELEVANCE:** Half of all children with high-risk neuroblastoma experience relapse or have refractory disease. These children lack effective options, and few survive beyond 5 years. The BCL2 inhibitor venetoclax, combined with cyclophosphamide/topotecan, has clinical activity in relapsed and refractory neuroblastoma, however, is limited by treatment-related toxicity. Combination drug screening in patient-derived models identified approved drugs synergistic with venetoclax, including the Aurora kinase A inhibitor alisertib. We assessed venetoclax-alisertib in 22 diverse neuroblastoma PDX models using an n=1 clinical trial-like design. Venetoclax-alisertib induced objective response in all models and was particularly effective against *MYCN*-amplified disease (n=12), doubling median survival time compared to cyclophosphamide/topotecan, and outperforming venetoclax/cyclophosphamide/topotecan. Efficacy was maintained with discontinuous schedules, minimizing hematological toxicity without substantially compromising activity. Venetoclax-alisertib was strikingly effective with anti-GD2 immunotherapy, allowing long-term survival of PDX-engrafted animals. Our findings support advancement of dual BCL2-Aurora kinase A inhibition to clinical trial for neuroblastoma with or without anti-GD2 immunotherapy.

## INTRODUCTION

Approximately 50% of children diagnosed with neuroblastoma have high-risk disease based on age at presentation, metastases and/or unfavorable genetics (e.g. *MYCN* amplification) (1). Despite the introduction of anti-GD2 directed immunotherapy in frontline therapy around 50% still experience relapse or have refractory disease, with few surviving beyond 5 years (2). Current treatments for high-risk NB (HR-NB) rely heavily on conventional cytotoxic chemotherapy, with most survivors experiencing long-term adverse effects attributable to their treatment (3, 4). With the limited success of the current therapies for HR-NB patients, there is a critical need to develop safer and more effective treatments.

While targeted agents have revolutionized the treatment of some cancers, their application in HR-NB is limited by the paucity of recurrent druggable mutations or fusions, with a rare exception being the activity of ALK inhibitors in ALK mutant HR-NB, an aberration present in ∼9% of HR-NB at diagnosis (5). HR-NB outcomes have been improved by the introduction of anti-GD2 immunotherapy (e.g. dinutuximab) in the post-consolidation phase, which has increased event-free survival (6) and has recently been shown to increase response rates at induction (7). As anti-GD2 becomes more widely used at induction, new chemoimmunotherapy combinations that augment this activity will be needed, as will new targeted agent combinations for patients who are resistant to upfront anti-GD2 therapies (8, 9).

The B-cell lymphoma 2 (BCL2) family of anti-apoptotic proteins, which regulate the intrinsic apoptosis pathway, have emerged as promising drug targets for HR-NB. Of particular interest is BCL2 itself, which has high relative expression in these tumors (10–12). Venetoclax, a selective BCL2 inhibitor, binds to BCL2 and displaces the pro-apoptotic protein BCL2-interacting mediator (Bim), which subsequently activates the effector proteins BAX and BAK, causing mitochondrial damage and cytochrome c release, leading to initiation of apoptosis (13). Venetoclax is highly tolerable and has been approved for clinical use in chronic lymphocytic leukemia (CLL) and acute myeloid leukemia, with a 70% response rate in relapsed CLL (14). Recent data indicates that, as with adult blood cancers (15–17), venetoclax is most effective against HR-NB when used in combination. Response of a venetoclax sensitive cell line xenograft was shown to be further improved by combination with cyclophosphamide (18), and *MYCN*-amplified neuroblastoma PDX models are more sensitive to venetoclax when combined with cyclophosphamide and topotecan than monotherapy (10). A recent phase 1 study in children and young adults with relapsed or refractory solid tumors, including HR-NB (NCT03236857) indicated an objective response rate of 31% for venetoclax with cyclophosphamide and topotecan in HR-NB; however, with a need for discontinuous dosing to combat neutropenia and thrombocytopenia (19, 20), corroborating a small case series reporting similar findings (19, 20). Furthermore, venetoclax in combination with other targeted agents has been studied in preclinical models, harboring specific molecular aberrations. For example, venetoclax in combination with the MDM2 inhibitor idasanutlin was effective in HR-NB models with wild-type *TP53* and BCL2 dependency (21, 22) and its combination with the MEK inhibitor trametinib was effective in models with RAS-MAPK aberrations (23), while the combination with alisertib was active in models with *MYCN* amplification (24). Despite promising results, the generalizability of their antitumor effects in broader HR-NB models remains unexplored. Combination of venetoclax with inhibitors of another pro-apoptotic member of BCL2 protein family, MCL1, has also been studied (10, 25), however with limited preclinical activity.

Here, we aimed to uncover novel drug combinations with venetoclax that are effective across neuroblastoma subtypes and may be better tolerated than its combination with cyclophosphamide and topotecan. We performed unbiased high-throughput combination screening with venetoclax and tested several synergistic combinations in a subset of HR-NB PDX models. Using a clinical trial-like n=1 study, we showed that the combination of venetoclax-alisertib was broadly effective, inducing objective response across a wide range of tumors, with strongest activity in tumors with *MYCN* amplification, where it outperformed the triple combination of venetoclax-cyclophosphamide-topotecan. We also report that the effect of venetoclax-alisertib was maintained with less frequent dosing and was strikingly effective in combination with anti-GD2 immunotherapy.

## MATERIALS AND METHODS

### Ethical approvals and consent

All animal experiments were approved by the University of New South Wales Animal Care and Ethics Committee (ACEC 20/119B, 22/145B, 23/79B) and conducted in accordance with the Animal Research Act (New South Wales) and the Australian code for the care and use of animals for scientific purposes. De-identified models were developed from consented patients, approved by the Sydney Children’s Hospital Network Human Research Ethics Committee (2020/ETH01981).

### Patient-derived xenograft models

HR-NB PDX models were established at Children’s Cancer Institute from patients enrolled in Australia’s pediatric personalized medicine trial (PRISM, NCT03336931) or consented for biobanking for research through Sydney Children’s Hospital Network (26). Additional models were obtained from the Children’s Oncology Group repository (Lubbock, Texas, USA). For expansion, dissociated tumor samples were suspended in RPMI/Matrigel mix (1:1, v/v) and inoculated subcutaneously into immunodeficient female NOD.Cg-*Prkdc^scid^Il2rg^tm1Wjl^*/SzJ (NSG) mice from Australian Bioresources (Moss Vale, NSW, Australia) or OZgene ARC (Perth, WA, Australia). At 1000mm^3^ tumor (volume = length × width × height/2 measured by Vernier caliper), mice were euthanized and tumor tissue snap frozen, dissociated into single cells, or cryopreserved.

### Co-immunoprecipitation and western blot

Tumor pieces were homogenized in 1mL 2% CHAPS buffer with protease inhibitors (Thermo Scientific) using a TissueRuptor homogenizer (Qiagen), then lysed overnight at 4°C followed by centrifugation at maximum speed for 20 min. Lysates were precleared of non-specific proteins by incubation with CHAPS-washed Protein A/G Agarose beads for 1.5h (4°C). Equal protein amounts were incubated for 1.5h with either the BCL-2 antibody (Cell Signaling) or a FLAG Tag antibody (Cell Signaling) then exposed to protein A-agarose beads (Roche) overnight at 4°C to capture protein complexes. Co-immunoprecipitated proteins were isolated following washing and centrifugation cycles (2 min, 8000 rpm, 4°C). Equal amounts of protein were diluted in 5× Laemmli buffer with DTT, boiled (95°C, 5 min), separated on 4–20% mini-PROTEAN TGX gels (Bio-Rad), and transferred to nitrocellulose membrane (Amersham). Membranes were blocked with 5% skim milk powder in TBS-T for 1h, incubated overnight (4°C) with primary anti-BCL2 or anti-Bim antibodies (Cell Signaling, 1:1000 dilution; rabbit mAb in TBS-T with 5% w/v skim milk), then for 2h with secondary antibody (1:5000 dilution; anti-rabbit HRP conjugate in TBS-T), washing 3×5 min in TBS-T between steps. Protein bands were visualized by incubation in Clarity ECL substrate for 2 min and detection with chemiluminescent and colorimetric imaging using a ChemiDoc MP Imaging System (Bio-Rad). Bands were quantified using ImageLab (Bio-Rad). Anti-Actin (rabbit polyclonal, 1:5000; in TBS-T) or anti-GAPDH (mouse polyclonal 1:10000; in TBS-T) antibodies were used as loading controls.

### Immunohistochemistry

Tissue microarrays (TMA) containing 34 NB PDX models (2–3 cores/model) and normal tissue controls were constructed at NSW Health Statewide Biobank from FFPE blocks. TMA blocks were cut into 4 µM sections for hematoxylin and eosin (H&E) and immunohistochemistry (IHC) staining (Garvan Institute of Medical Research Histopathology Services). IHC included antigen retrieval at 100°C for 30 min in Bond Epitope Retrieval solution 2 (Leica Biosystems #AR9640), and 60 min incubation with primary antibodies directed against PHOX2B (Abcam ab183741; 1:1000 dilution) and BCL2 (ThermoFisher Scientific MA5-11757; 1:300 dilution) using the Leica Bond Polymer Refine system (Leica Biosystems). Staining was scored by a pediatric pathologist (AJG). Positive nuclear staining for PHOX2B confirmed the diagnosis of neuroblastoma in each core. BCL2 staining intensity was designated 0 (negative), 1 (weak), 2 (moderate), or 3 (strong), and proportion of positive tumor cells designated 0 (0%), 1 (<10%), 2 (10–50%), or 3 (>50%). The final score was intensity × proportion, range 0–9.

### WGS data analysis

Library preparation and whole genome sequencing analysis was conducted at the Australian Genome Research Facility (Melbourne, Australia), using the Illumina Novoseq X platform with a paired-end read length of 150 bases. Methods have been previously detailed (27). *MYCN* amplification (>10 copies) was called according to INRG criteria (28).

### Ex vivo high-throughput combination drug screening

Freshly dissociated PDX tumor cells were seeded in 384-well plates (Greiner) using Multidrop Combi (Thermo) reagent dispensers (5,000 cells per well) and incubated for 72h. A Tecan HP D300 Digital dispenser was used to add DMSO vehicle or venetoclax (Adooq Bioscience) at the IC_30_ concentration for each PDX model, followed by a library of 111–126 drugs approved or in clinical or pre-clinical development for childhood cancer, dispensed in duplicate as 10-fold serial dilutions 0.5nM–5µM using a Microlab® STAR^TM^ liquid handling robot (Hamilton). After 72h treatment, viability was measured using CellTiter Glo 2D (Promega). Raw values were converted into percentage viability using the formula: ([readout value drug – average readout positive controls]/[average readout negative controls – average readout positive controls] × 100) and used to generate dose-response curves using a four-parameter logistic function and to calculate the area under the dose–response curve (AUC) and half maximal inhibitory concentrations (IC_50_) values. Synergy was determined by average Bliss independence score using SynergyFinder (https://synergyfinder.fimm.fi) (29).

### In vivo efficacy studies

Dissociated PDX tumor cells (1–2×10^6^ cells/animal in equal volume of RPMI and Matrigel) were subcutaneously engrafted into 6-week-old female NSG mice or for experiments including anti-GD2 immunotherapy, BALB/c nude mice (OZgene ARC) and randomized into treatment groups of four mice/treatment group/model for conventional experiments, or one mouse/treatment group/model for single-mouse trial format (SMT) (30). Treatment commenced at tumor ≥100 mm^3^. Schedules included 100 mg/kg PO venetoclax once daily for 5 consecutive days for 4 weeks in 10% ethanol/30% polyethylene glycol/60% phosal, 10.4 mg/kg PO alisertib (Puma Biotechnology) twice daily for 5 consecutive days for 4 weeks in 10% 2-hydroxypropyl-β-cyclodextrin, 1% NaHCO_3_ in water, 35 mg/kg IP vorinostat (Selleck Chemicals) once daily for 5 consecutive days for 4 weeks in 2% DMSO/30% PEG300/5% Tween80 in water, 0.5 mg/kg, IP vincristine (Selleck Chemicals) once weekly for 4 weeks in saline, and 6 mg/kg Etoposide (Selleck Chemicals), 20 mg/kg cyclophosphamide (Baxter Healthcare) and 0.5 mg/kg topotecan (Sandoz) each IP once daily for 5 days and repeated at day 21 in saline. Alternative venetoclax and alisertib schedules were as described in Results, with doses and formulation unchanged. Hematology parameters were measured at day 21 and day 42 after start of treatment using Auto Hematology Analyzer (Mindray BC-5000). For chemoimmunotherapy, tumor cell GD2 expression was confirmed by flow cytometry using FITC anti-human GD2 (BD, 1:1000) and corresponding FITC anti-human IgG isotype (BD, 1:1000). Venetoclax, alisertib, cyclophosphamide and topotecan were administered as above. 2 mg/kg Irinotecan (Accord Healthcare) and 5 mg/kg temozolomide (Sigma-Aldrich) were delivered IP, once daily for 5 days and repeated at day 21 in saline. Mouse IgG2a isotype (Bio-X-Cell) and anti-GD2 [clone 14G2a] (Bio-X-Cell) were given IP in 2 doses of 15 µg on days 1 and 5. Tumor volume was measured at least twice weekly. Mice were euthanized when the tumor reached 1000mm^3^. Study endpoint (event) was defined as quadrupling of tumor size from the start of treatment. Time to event was assessed using Kaplan-Meier survival curves which were compared using the Mantel-Cox log-rank test. Response to treatment was determined according to objective response measures set by Pediatric In Vivo Testing consortium (PIVOT) and categorized as progressive disease 1 or 2 (PD1 or PD2), stable disease (SD), partial response (PR), complete response (CR), and maintained complete response (MCR) (31). Event free survival (EFS) T/C value is determined by the ratio of median time to event of the drug-treated group (T) over the control group (C).

### Statistical analysis

Analyses were performed in GraphPad Prism 11.0.0 (GraphPad Software). Event-free survival between treatment and control was compared using Kaplan-Meier analysis followed by Mantel-Cox test: * P<0.05, ** P<0.01, ***P<0.001. Correlation plots were generated by linear regression analysis.

### Data availability

The genomic data generated in this study are publicly available from the European Genome-phenome Archive https://ega-archive.org/ under accession number EGAS00001008494. Previously published genomic datasets are available under accession number EGAS00001004572, EGAS00001008220, EGAS00001004905, EGAS00001005811, EGAS00001007029. Genomic data for PDX models from Children’s Oncology Group is available on PedCBioPortal database https://pedcbioportal.kidsfirstdrc.org/. All other raw data generated in this study are available upon request from the corresponding author.

## RESULTS

### BCL2 is broadly expressed across models of high-risk neuroblastoma, and inhibition sensitizes to chemotherapy

We first examined BCL2 protein expression by IHC across a panel of 34 HR-NB PDX models. BCL2 staining was non-nuclear. While a broad range of expression levels were observed, as reflected by TMA protein expression score, 33 of 34 models expressed BCL2 (Figure 1A). High BCL2 expressing models (e.g. zccs373) typically showed strong diffuse staining, while moderate BCL2 expressing models (e.g. zccs51) had less uniform staining and low BCL2 expressing models (e.g. zccs242) showed only occasional positive cells (Figure 1B, Supplementary Figure S1). The breadth of BCL2 protein expression was largely reproduced by western blotting (n=21; Supplementary Figure S2).

**Figure 1.**
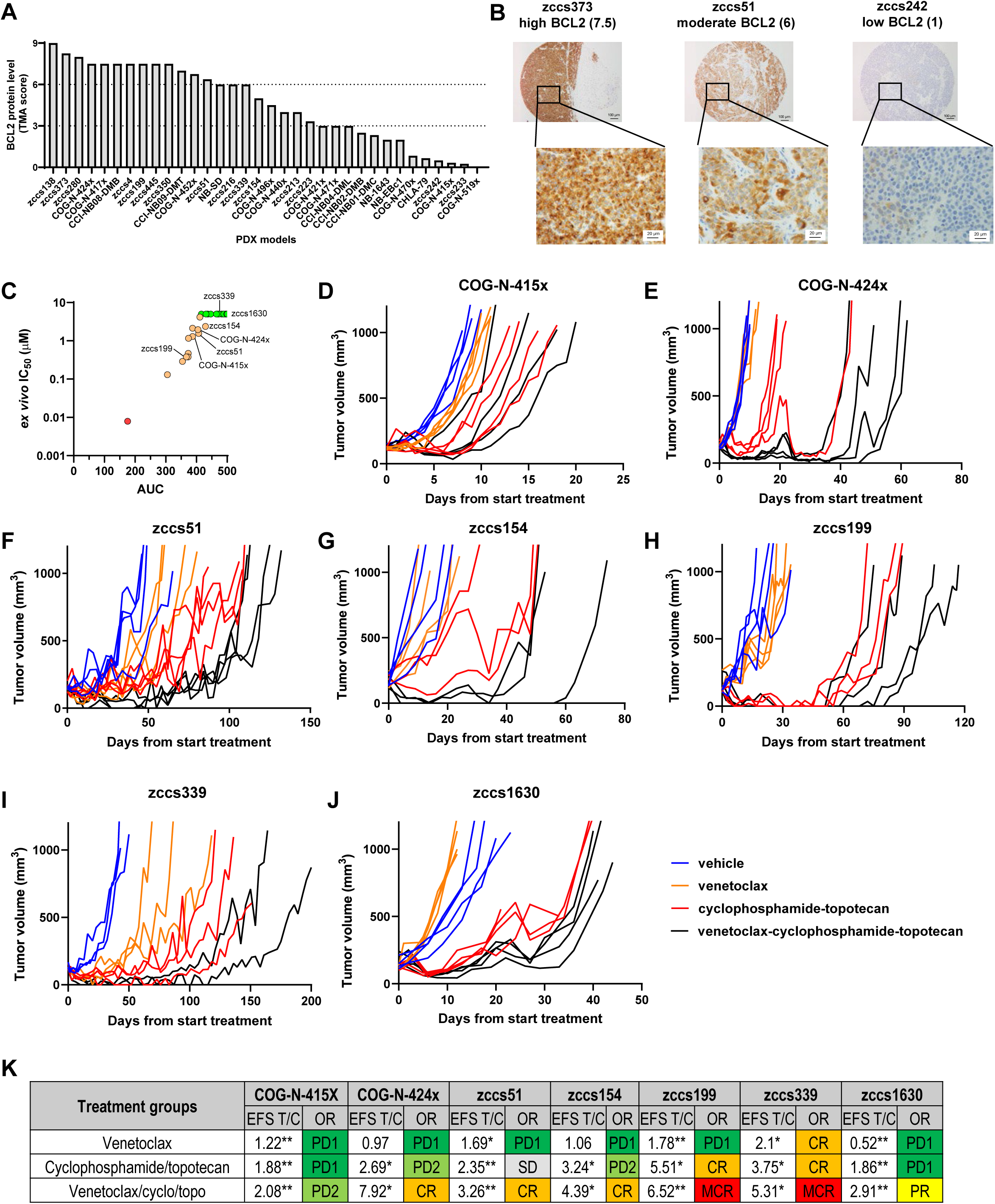
BCL2 expression and venetoclax response in HR-NB PDX models. (**A**) IHC assessment of BCL2 protein levels in HR-NB PDX models (n=34) based on scoring of TMA cores (0–9 based on intensity and positively stained proportion, averaged over multiple cores). (**B**) TMA cores stained for BCL2, illustrating high-, moderate-, and low-expression. Images shown at 100× and 600× magnification, with TMA score indicated. **C)** *In vitro* sensitivity of venetoclax across a panel of HR-NB PDX model based on IC_50_ value and AUC and colored by sensitivity: red = sensitive (IC_50_ <0.01µM), orange = intermediate (1µM<IC_50_<5µM), green = insensitive (IC_50_>5µM). (**D–J**) Tumor growth curves of HR-NB PDX models in NSG mice treated with vehicle, venetoclax, cyclophosphamide-topotecan or combination. (**K**) Magnitude of drug responses were indicated by EFS T/C value and objective response criteria. Differences between treatment arms and vehicle group were analyzed using Log-rank test \**P*<0.05, \*\**P*<0.01.

We next analyzed venetoclax sensitivity in HR-NB PDX models *ex vivo* (n=30). Of these, one model was sensitive at low nanomolar concentration (IC_50_ 8 nM; AUC 171). An additional twelve models had a measurable response to venetoclax at higher doses (IC_50_ 0.13 µM–4.2 µM; AUC 305–429), while the remaining 17 models were resistant to venetoclax at 5 µM (Figure 1C). We then assessed *in vivo* responses to venetoclax (100 mg/kg PO) with and without cyclophosphamide-topotecan in a subset of seven PDX models in NSG mice. Most models were resistant to venetoclax monotherapy with mice experiencing progressive disease (PD1) while for one model, zccs339, mice achieved complete response (CR; Figure 1D–K).

This response was not predicted by *ex vivo* venetoclax sensitivity (Figure 1C). Responses to cyclophosphamide-topotecan ranged from PD to CR, however in all but one model (COG-N-415, with very low BCL2 expression), combination with venetoclax improved objective response beyond that observed for cyclophosphamide-topotecan alone (Figure 1K), supporting the application of venetoclax as a chemosensitizer in combination therapy.

### BCL2 and BCL2-Bim complex levels are not definitive predictors of venetoclax sensitivity

To determine whether BCL2-Bim complex levels could predict venetoclax sensitivity in PDX models, we first performed BCL2 pulldown and western blotting for co-immunoprecipitated Bim and BCL2. Both proteins were quantitated after normalization to loading controls. The neuroblastoma cell line SMS-KCNR, previously demonstrated to have high BCL2-Bim complex levels, was used as an internal reference (11). The levels of co-immunoprecipitated BCL2-Bim complex varied substantially across the PDX model panel (n=33; Figure 2A), with a summary heatmap (relative to SMS-KCNR for each measure) shown in Figure 2B. Levels of complexed Bim and BCL2 were correlated with responses to venetoclax measured *ex vivo* (AUC values; Figure 1C) and *in vivo* (EFS T/C values; Figure 1K, Supplementary Figure S3 for zccs223, and Figure 4D for COG-N-440x and zccs373). We found that *ex vivo* AUC values were not closely correlated with complexed Bim or BCL2 levels (P=0.056 and P=0.04 respectively; Figure 2C, D). While complexed Bim levels were significantly correlated with venetoclax EFS T/C values (P<0.0001; Figure 2E), this response was heavily influenced by a single model with exceptional response (zccs373) and very high BCL2 expression which we subsequently confirmed as an outlier using Grubbs analysis (Alpha = 0.05, G score = 2.626, P value excluding zccs373 = 0.033). BCL2 protein levels were not correlated with venetoclax EFS T/C values (P = 0.885; Figure 2F). Taken together, our data does not support BCL2-Bim complex levels or BCL2 protein levels as definitive indicators for venetoclax sensitivity.

**Figure 2.**
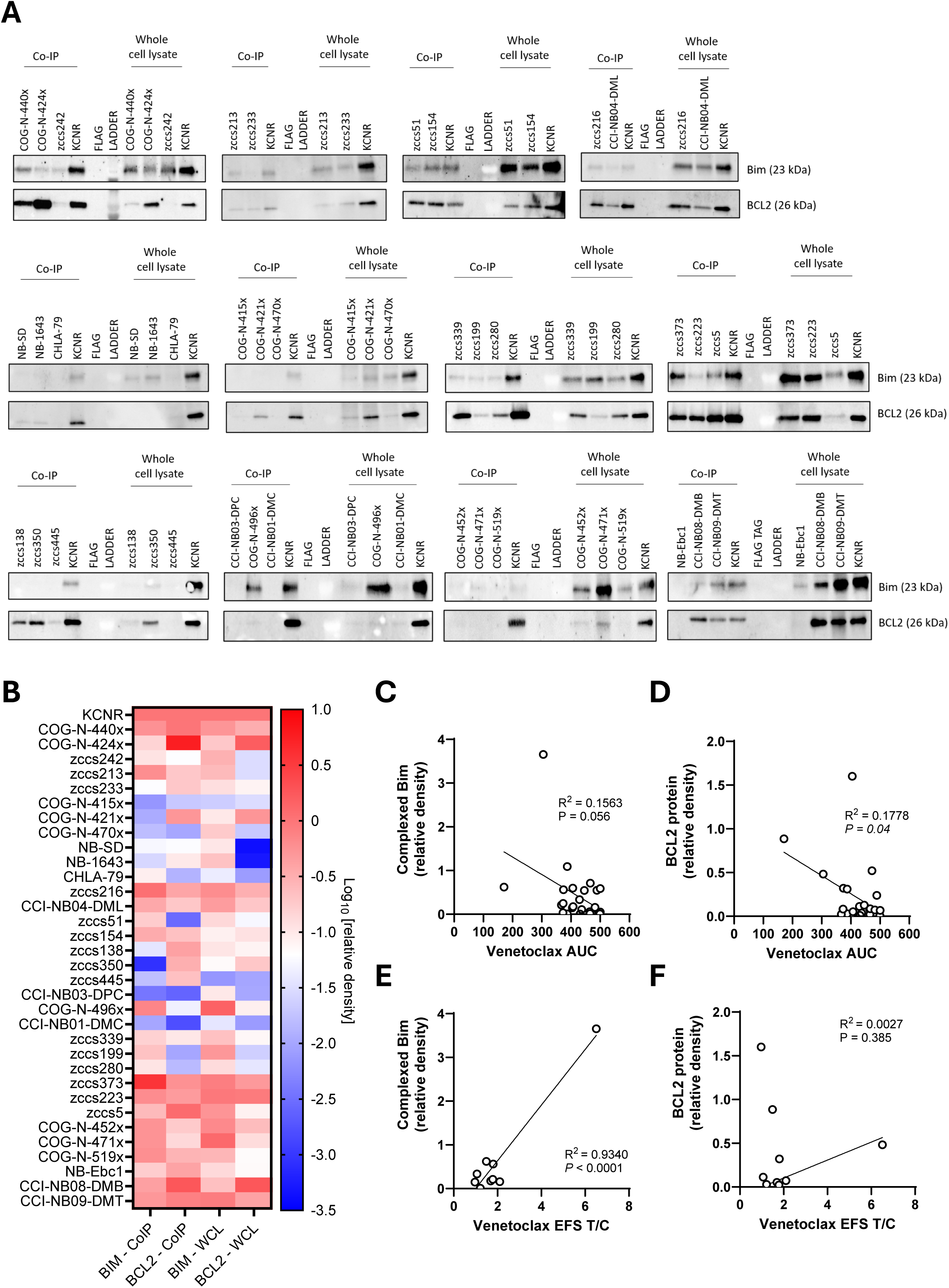
Complexed Bim-BCL2 correlated with venetoclax sensitivity *in vivo* but not *ex vivo* in HR-NB PDX models. (**A**) Western blot images from Bim-BCL2 co-IP and whole cell lysate (WCL) of HR-NB PDX models. SMS-KCNR cell lines were included as a positive control for Bim-BCL2 complex. FLAGtag was a negative control antibody for co-IP. (**B**) Heatmap of the Log value of relative optical density on BCL2 and Bim-BCL2 blots. Co-IP and WCL samples were normalized to their respective KCNR reference, and to GAPDH or beta-actin loading controls of respective blot. (**C**) Correlation of *ex vivo* venetoclax AUC with complexed Bim protein levels (R^2^ = 0.1563. *N.S.*: not significant) and (**D**) BCL2 protein levels (R^2^ = 0.1778. P=0.04). (**E**) Correlation of median EFS T/C of venetoclax monotherapy *in vivo* with complexed Bim protein levels (R^2^ = 0.9340, P<0.0001) and (**F**) BCL2 protein levels (R^2^ = 0.0027, *N.S.*: not significant).

**Figure 3.**
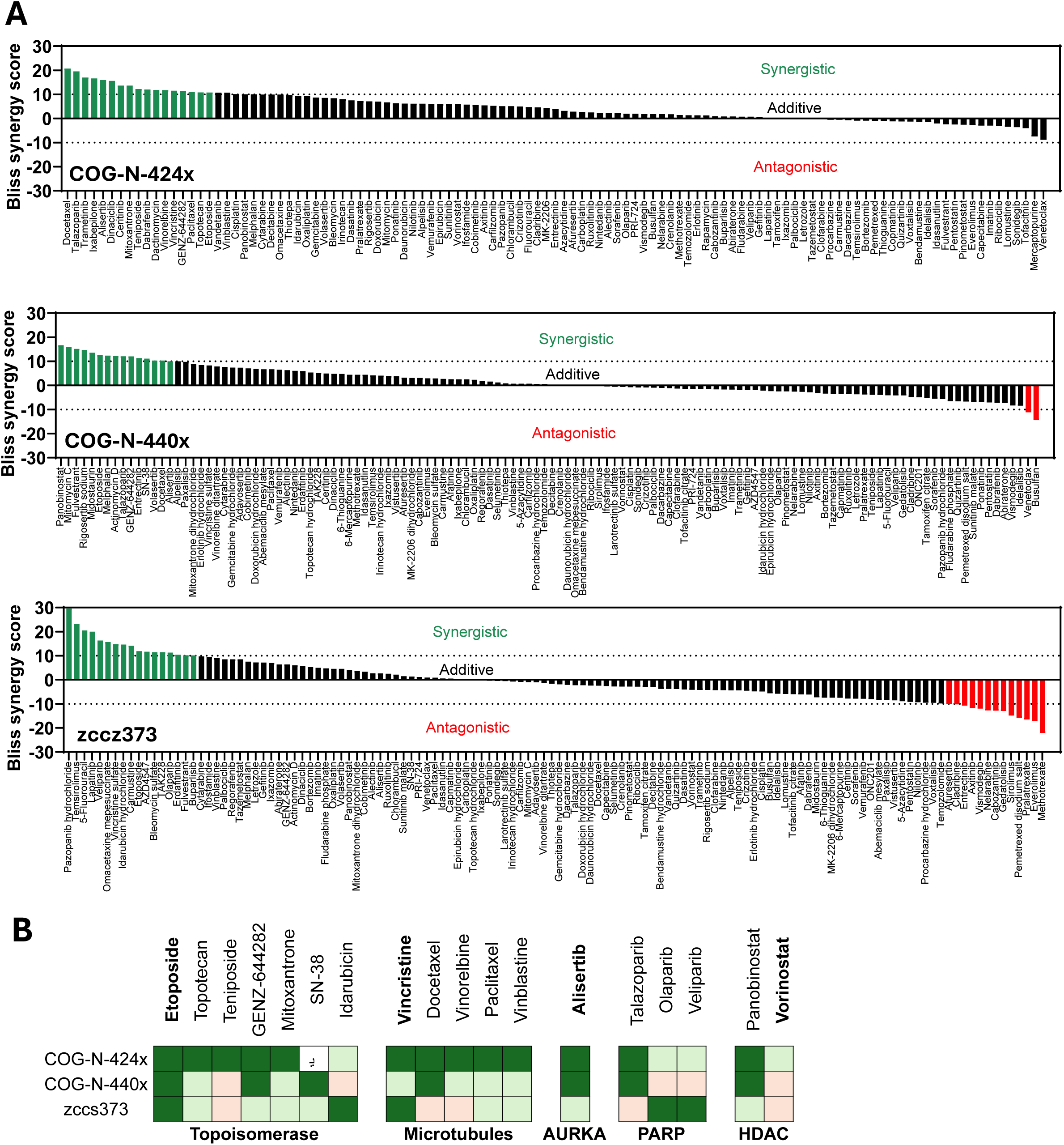
High-throughput drug screening identifies drugs synergistic with venetoclax in HR-PDX models. (A) Combination screens were conducted in three models: COG-N-424x, COG-N-440x, and zccs373 with venetoclax (IC30) and library drugs at 10-fold dilution from 5 μM to 0.5 nM. Drug synergistic with venetoclax shown in green (Bliss Score ≥ 10) and those antagonistic in red (Bliss Score ≤ –10). (B) Recurrent synergistic drugs or drug classes. Synergistic drugs shown in dark green and additive drugs shown in light green (0 < Bliss Score < 10) or pink (−10 < Bliss score ≤ 0). Vorinostat include for comparison. “–” = datapoint failed quality control.

**Figure 4.**
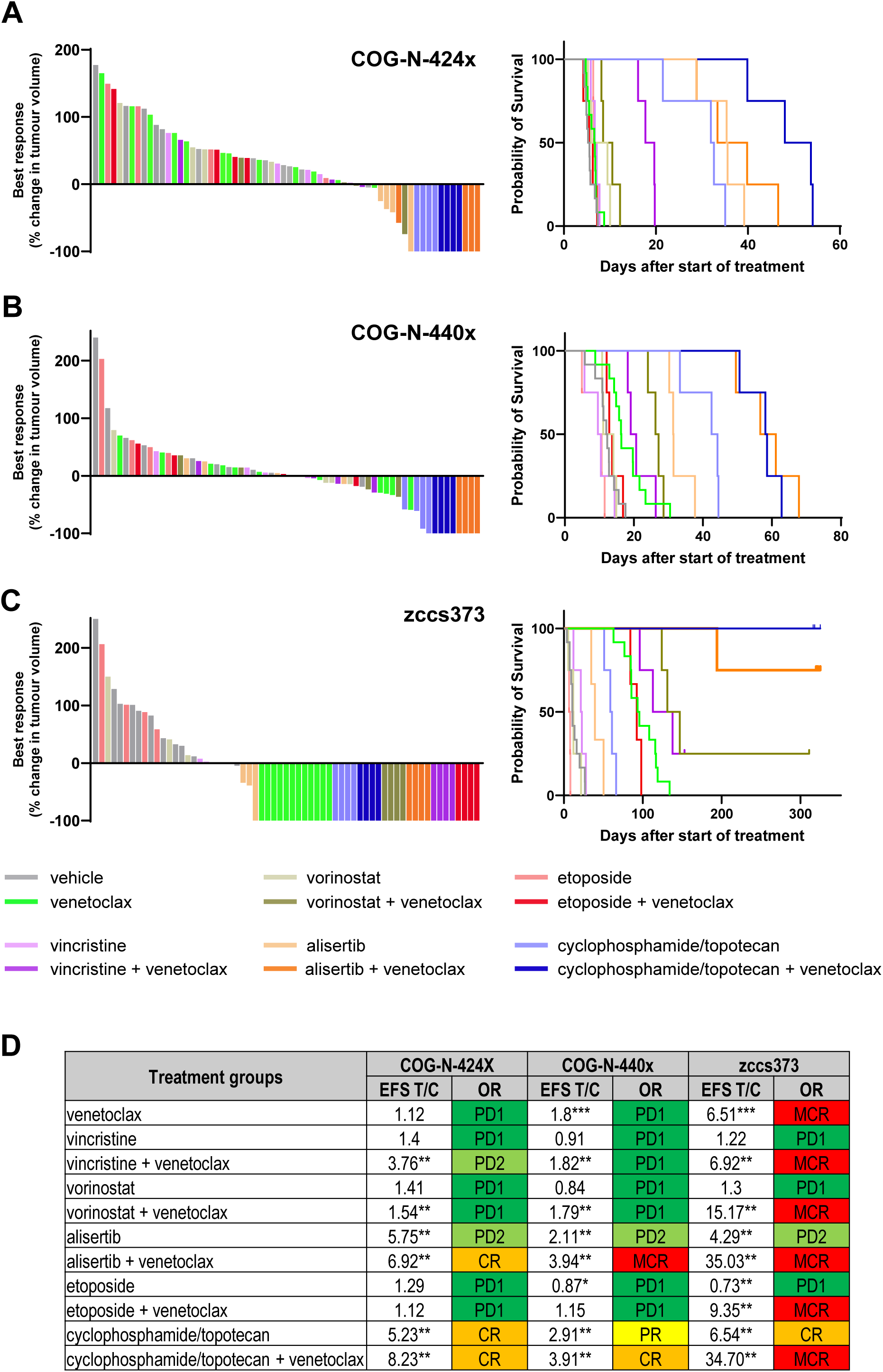
*In vivo* studies indicate venetoclax combination with alisertib as the best performing combination in HR-NB PDX models. **(A)** Best response waterfall plot and Kaplan-Meier survival curve for COG-N-424x, **(B)** COG-N-440x, and **(C)** zccs373 in NSG mice. Colored bars in waterfall plot indicate the best response obtained for individual mice included in different treatment arms. **(D)** Magnitude of drug responses were indicated by EFS T/C value and objective response criteria. Differences between treatment arms and vehicle group were analyzed using Log-rank test \**P*<0.05, \*\**P*<0.01, ***P<0.001.

### High-throughput combination screening identifies synergistic venetoclax combinations

To identify agents that are synergistic with venetoclax, we performed unbiased *ex vivo* high-throughput drug screening with a library of anticancer drugs containing 111–126 anticancer drugs, enriched for those approved or in clinical or pre-clinical development for childhood cancer. Three HR-NB PDX models were selected for the screening based on the relative levels of BCL2-Bim complex in each model, zccs373 (high), COG-N-440x (intermediate), and COG-N-424x (low). Venetoclax was at the IC_30_ for each model. Combinations were ranked by Bliss Synergy score across the dose range, and designated as synergistic with a Bliss score ≥ 10, additive with a Bliss score between -10 and 10, and antagonistic with a Bliss score ≤ –10. Venetoclax was synergistic with 25 drugs in COG-N-424x, 15 drugs in COG-N-440x, and 17 drugs in the zccs373 (Figure 3A). Recurrent synergistic drugs or drug classes included inhibitors of DNA topoisomerase, microtubules, AURKA, PARP, and HDAC (Figure 3B).

### Venetoclax-alisertib combination outperformed other combinations in vivo

Recurrent synergistic combinations identified by drug screening were assessed *in vivo* using the same three PDX models as the *ex vivo* screen, each engrafted in NSG mice. Drugs assessed included the standard-of-care agents etoposide and vincristine, and the AURKA inhibitor alisertib. Despite additivity rather than synergy, the pan-HDAC inhibitor vorinostat was included as an alternative to panobinostat based on more extensive clinical evidence for therapeutic efficacy in this disease. Standard-of-care cyclophosphamide-topotecan and venetoclax-cyclophosphamide-topotecan were included for comparison. Waterfall plots for best response for each animal, along with survival curves (EFS) are shown in Figure 4A–C, while all tumor growth curves are shown in Supplementary Figure S4. Extension of survival (EFS T/C) and objective responses for each combination are shown in Figure 4D. COG-N-424x and COG-N-440x were resistant to venetoclax monotherapy (PD1), while zccs373 was highly sensitive, achieving MCR (EFS T/C = 6.51, *P* < 0.001). All three models responded to cyclophosphamide-topotecan (PR–CR) and to cyclophosphamide-topotecan-venetoclax (CR– MCR; Figure 4A–D, Supplementary Figure S4). Of the combinations identified by *ex vivo* screening, the most effective *in vivo* was venetoclax-alisertib which achieved CR in COG-N-424x (EFS T/C 6.92; *P* = 0.0067), MCR in COG-N-440x (EFS T/C 3.94; *P* = 0.0067) and MCR in zccs373 (EFS T/C 35.03; *P* = 0.0067), comparable to the triple combination of venetoclax-cyclophosphamide-topotecan. Alisertib monotherapy did not induce objective response in these models (PD2), indicating that strong synergy between venetoclax and alisertib could be achieved even in the PDX models that were insensitive to venetoclax or alisertib monotherapy. The effectiveness of the other venetoclax combinations was more limited. No other combination reached objective response in the COG-N-424x and COG-N-440x models, and zccs373 achieved MCR with venetoclax alone. However, extension of survival (EFS T/C) was increased with vincristine-venetoclax in COG-424x, and with both vorinostat-venetoclax and etoposide-venetoclax in zcc373. Mice tolerated all combination treatments and maintained stable body weights (Supplementary Figure S5).

### Venetoclax-alisertib is broadly effective against HR-NB, with enhanced activity against MYCN-amplified tumors

As venetoclax-alisertib induced CR or MCR in all three PDX models, we extended assessment to a larger cohort more representative of the heterogeneity of high-risk neuroblastoma, using a clinical-trial-like single mouse study design. The cohort was established from 20 patients (11 males and 9 females), age at diagnosis 0.5–13.1 years, including paired diagnosis-relapsed models from two patients, total 22 models. Diverse molecular features were represented, including aberrations in *MYCN*, *ALK*, RAS-MAPK pathway genes, *TP53*, *TERT* and *ATRX*. Each model was represented by one NSG mouse per treatment arm, comparing vehicle, venetoclax-alisertib, cyclophosphamide-topotecan (standard-of-care chemotherapy comparator), and venetoclax-cyclophosphamide-topotecan (combination from phase 1 trial). To include data from the three models already assessed (COG-N-424x, COG-N-440x, and zccs373; Figure 4), median responses from 4 animals per group were plotted as single data point for each treatment group. Venetoclax-alisertib treatment achieved objective response in all models tested, with PR in two models and CR–MCR in all other models (Figure 5A, B, Supplementary Figure S6). Across all models, EFS was substantially extended by venetoclax-alisertib compared to vehicle (EFS T/C = 9.62, P < 0.0001) (Figure 5C). Venetoclax-alisertib outperformed standard-of-care cyclophosphamide-topotecan (P = 0.0180; EFS T/C = 6.14) and was comparable to the triple combination of venetoclax-cyclophosphamide-topotecan (P=0.535; EFS T/C = 8.22). Venetoclax-alisertib was particularly effective in models of *MYCN*-amplified disease (EFS T/C = 13.05, Figure 5D) where extension of survival exceeded that of *MYCN* non-amplified models (Supplementary Figure S7A), despite *MYCN*-amplified models tending to be less sensitive to cyclophosphamide-topotecan than their non-amplified counterparts (median EFS = 44.34 *vs* 80.18, P = 0.1375 n.s.; Supplementary Figure S7B). In contrast, in *MYCN* non-amplified models, venetoclax-alisertib response was comparable to that of cyclophosphamide-topotecan (Figure 5E).

**Figure 5.**
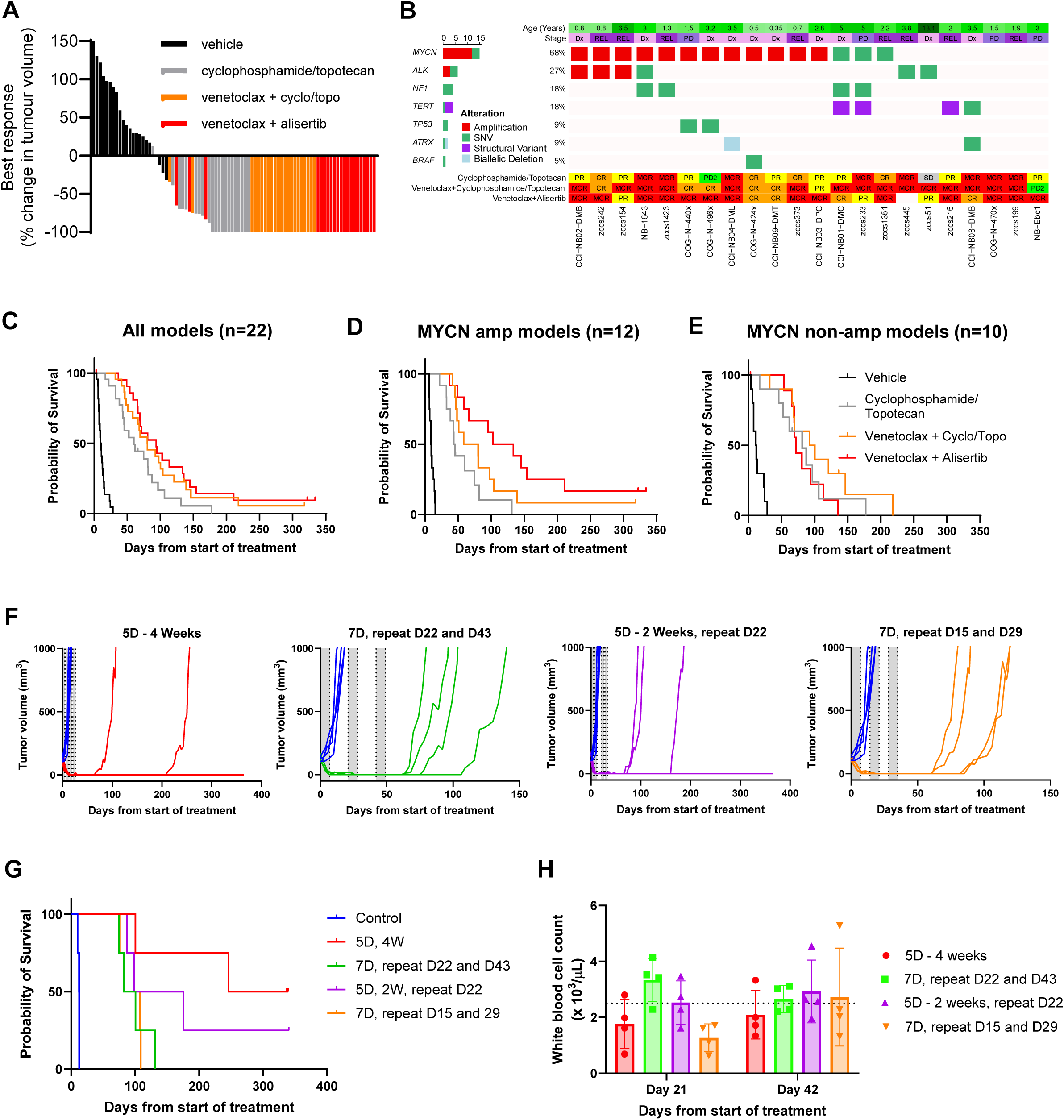
Venetoclax-alisertib is highly effective in a wide range of HR-NB PDX models and maintains efficacy in discontinued dosing. (**A**) Best response waterfall plot and from a clinical trial-like (n=1) study in 22 HR-NB PDX models. (**B**) Demographic and molecular characteristics of the HR-NB PDX models and objective response criteria following venetoclax combination treatments. Kaplan-Meier survival curves for all models (**C**), *MYCN*-amplified models (**D**) and *MYCN* non-amplified models (**E**). (**F**) Tumor growth curves for COG-N-440x engrafted mice treated with vehicle (blue) or alternative venetoclax-alisertib schedules, including for 5 days for 4 weeks (red), 7 days with repeat at day 22 and day 43 (green), 5 days for 2 weeks with repeat at day 22 (purple), and 7 days with repeat at day 15 and day 29 (orange). Each line represents one mouse (n=4/group). Shaded areas represent the treatment period. (**G**) Kaplan-Meier survival curves for mice treated with different schedules of venetoclax-alisertib combination. (**H**) White blood cell counts of mice treated with different schedules of venetoclax-alisertib combination. Dotted line represents average white blood cell level in untreated NSG mice.

### Discontinuous dosing minimizes hematological toxicity without substantially compromising activity

Clinical experience with both venetoclax with chemotherapy (20) and alisertib (32) indicates that discontinuous treatment strategies are required for minimizing hematopoietic toxicity. To investigate whether discontinuous dosing of venetoclax-alisertib compromised activity, we compared our standard 5d/week for 4 weeks dosing schedule to three alternative schedules with longer recovery time, namely: 7d/week repeat cycle at d22 and d43, 5d/week for 2 weeks repeat cycle at d22, and 7d/week repeat cycle at d15 and d29. Using the COG-N-440x model in NSG mice, we showed that MCR was achieved in all animals irrespective of schedule, and that disappearance of tumor was maintained for at least 60 days after start of treatment (Figure 5F). Longer-term tumor-free survival was, however, more likely to occur with the earlier intense treatment (Figure 5G). As expected, white blood cell count returned to normal in the two alternative schedules that were not on treatment at day 21 (Figure 5H), while at day 42, all groups were off treatment, hence white blood cell levels were normal. Platelet count returned to normal at both day 21 and day 42 only in the schedule with the longest treatment break (Supplementary Figure S8). Modified schedules can thus allow for recovery of the hematopoietic system without substantially compromising activity.

### Venetoclax-alisertib combination outperforms standard chemoimmunotherapy in neuroblastoma

We then compared the efficacy of venetoclax-alisertib with or without anti-GD2 antibody against standard chemoimmunotherapy using the COG-N-440x PDX model. GD2 expression was confirmed by flow cytometry (Supplementary Figure S9) before engraftment in a cohort of BALB/c mice, an immune-deficient strain that enables engraftment while maintaining capacity for antibody-dependent cytotoxicity (33). Isotype or anti-GD2 treatments did not reduce tumor size (PD) and only caused slight delay in tumor growth (Figure 6A-C, Supplementary Figure S10A). Isotype treatment with either irinotecan/temozolomide or cyclophosphamide/topotecan significantly prolonged mice survival (EFS T/C = 5.14 and 5.05, respectively). Combining the chemotherapies with anti-GD2 antibody decreased tumor size further to less than half of the original size (PR), in addition to prolongation of event-free survival (Figure 6A, C). Venetoclax-alisertib combination evidently outperformed the standard chemo-immunotherapies and caused disappearance of tumor for a prolonged period (MCR). Venetoclax-alisertib combination with anti-GD2 resulted in a longer event-free period (EFS T/C = >20.20) compared to its combination with isotype control (EFS T/C = 9.62) (Figure 6B, C). Mice tolerated venetoclax-alisertib-anti-GD2 treatments whereby their body weights were relatively stable throughout the treatment period (Supplementary Figure S10B).

**Figure 6.**
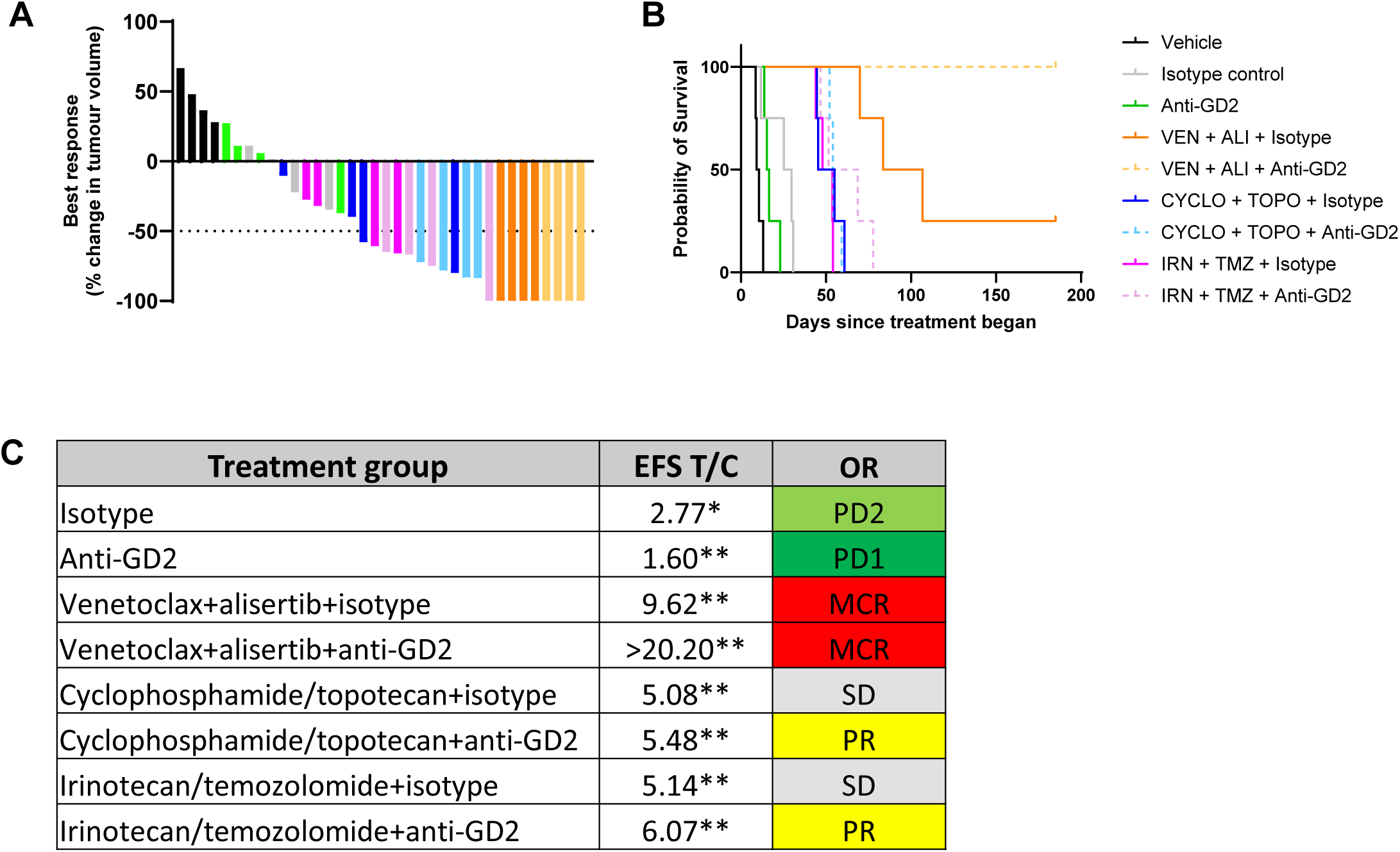
Venetoclax-alisertib outperforms standard chemoimmunotherapy and augments the effect of anti-GD2 therapy. (**A**) Best response waterfall plot and (**B**) Kaplan-Meier survival curve for COG-N-440x. Colored bars in waterfall plot indicate the best response obtained for individual mice included in different treatment arms. (**C**) Magnitude of drug responses were indicated by EFS T/C value and objective response criteria. Differences between treatment arms and vehicle group were analyzed using Log-rank test \**P*<0.05, \*\**P*<0.01.

## DISCUSSION

The identification and comprehensive evaluation of targeted therapy combinations is a high priority for the treatment of HR-NB therapy (9), a disease where treatment still relies heavily on conventional chemotherapy. A recent phase 1 study and a case series indicate activity for the combination of venetoclax-cyclophosphamide-topotecan in relapsed and refractory HR-NB patients (19, 20) (19, 20), but with substantial limitations imposed by treatment related toxicity. The potential for other combinations with BCL2 inhibitors, particularly targeted agents for treatment guided by molecular features, is evidenced by preclinical studies showing the activity of BCL2 inhibition with MDM2, MEK and AURKA inhibitors ((10), however the generalizability of these findings is underexplored. Ham *et al.* reported durable tumor regression in two *MYCN*-amplified cell line xenografts treated with venetoclax-alisertib, albeit at a higher dose of alisertib (30 mg/kg) than used here, with a non *MYCN*-amplified xenograft unresponsive (24). Here, we report that the combination of venetoclax with alisertib is highly active in a panel of 22 diverse HR-NB PDX models and that *MYCN* amplification is a biomarker of combination activity, with the most durable responses observed in models from this subset of patients with particularly poor outcomes. Mechanistically, the enhanced activity in *MYCN*-amplified neuroblastoma models is likely caused by disruption of AURKA-MYCN binding by alisertib resulting in MYCN degradation (34, 35). Downregulation of MCL1 via loss of p4E-BP1 and previously been proposed to contribute to the activity of this combination, (24), although the more restricted activity of dual BCL2-MCL1 inhibition that we have previously observed in xenograft models of HR-NB (25) suggests that this contribution in limited.

A unique element of our study was a clinical trial-like approach that assessed activity across HR-NB with a range of genetic features, including aberrations in *MYCN*, *ALK*, RAS-MAPK pathway genes, *TP53*, *TERT* and *ATRX*, improving the generalizability of our findings. Our study deliberately includes models from HR-NB subsets that, due to extended establishment times, are often under-represented in or absent from preclinical studies, such as older patients or those with *ATRX* mutant tumors. This approach also provides initial insights into the relative activity of venetoclax-alisertib compared to other venetoclax-based targeted agent combinations. We observed objective response (PR, CR, or MCR) for all models treated with venetoclax-alisertib, including in *NF1* mutant (n=4) and *TP53* wild-type (n=20). This contrasts with previous preclinical studies reporting stable disease in high BCL2-expressing and *NF1* mutant cells xenograft treated with venetoclax and trametinib (MEK inhibitor) (23), a lack of objective response to venetoclax with idasanutlin (MDM2 inhibitor) in the high BCL2 expressing, *TP53* wild-type PDX model COG-N-424x (22) (CR to venetoclax-alisertib), and to the less generalizable activity of venetoclax combined with the MCL1 inhibitor MIK665, which induced partial response in models zccs51 and COG-N-440x (25), which were PR and MCR respectively with venetoclax-alisertib. Therefore, while venetoclax-alisertib is most active in tumors with *MYCN* amplification, tumors with other targetable drivers may still benefit from this combination, and direct comparative studies will be required to ascertain superior combinations.

Notably, despite substantial BCL2 expression across the majority of our models, sensitivity to single agent venetoclax was rare, with *ex vivo* IC_50_ values typically above 1 µM and progressive disease *in vivo*. While various mechanisms of venetoclax resistance have been demonstrated, including upregulation of MCL1 (11), *BAX* mutation (36), *BCL2* mutation (37), and downregulation of PUMA and NOXA (38, 39), there are currently no definitive biomarkers of response to venetoclax for HR-NB. BCL2 protein levels and levels of BCL2-Bim complex have previously been reported to correlate with venetoclax sensitivity (11, 40), including in neuroblastoma cell lines. In our hands, neither BCL2 protein or BCL2-Bim complex levels were a robust predictor of venetoclax response, however we cannot exclude that more comprehensive approaches, such as BH3 profiling (41) could be more informative, as previously demonstrated for AML (42) (43).

While the activity of venetoclax-alisertib in our preclinical models is robust and compares favorably to both standard-of-care and other venetoclax combinations, tolerability is likely to present challenges for clinical use. Early phase trials of venetoclax-cyclophosphamide-topotecan (19, 20) and alisertib (32) required discontinuous treatment to manage neutropenia, potentially compromising activity. Here we showed that a clinically tolerable dose of alisertib in combination with venetoclax still induces MCR in preclinical models, even while incorporating 14-day breaks between treatment cycles, a schedule similar to that previously trialed for alisertib-irinotecan-temozolomide (44), increasing the feasibility of translation.

Incorporation of αGD2 immunotherapy into standard-of-care treatment has notably improved outcomes for HR-NB, and new chemoimmunotherapy combinations are an opportunity to build on this success (8) (9). Here we demonstrate highly durable responses and long-term tumor-free survival in a PDX model engrafted in BALB/c nude mice, with anti-GD2 antibody therapy administered at a dose equivalent to plasma concentration achievable in patients (under submission, preprint https://doi.org/10.21203/rs.3.rs-10274710/v1) (45). Our preclinical findings therefore support venetoclax-alisertib as a treatment strategy for patients refractory to immunotherapy, and as a potential chemoimmunotherapy combination for patients not previously treated with anti-GD2 antibody therapy.

In conclusion, we demonstrate that dual inhibition of BCL2 and AURKA is broadly efficacious in preclinical models of HR-NB, with enhanced activity against *MYCN*-amplified disease, and promising combinatorial activity as chemoimmunotherapy with anti-GD2 antibody. Discontinuous schedules to mitigate hematological toxicity appear feasible, reducing the barriers to clinical translation.

## ACKNOWLEDGEMENTS

Children’s Cancer Institute Australia is affiliated with the University of New South Wales and the Sydney Children’s Hospital Network. We acknowledge the ZERO Childhood Cancer Program Preclinical Team for providing HR-NB PDX samples and thank Dr. C. Patrick Reynolds and Children’s Oncology Group Childhood Cancer Repository (cccells.org) for providing additional models. We thank Dr. Angela Xie for assistance with validating GD2 expression by flow cytometry, and Ms. Crystal Mak, Ms. Jennifer Brand, and Ms. Caitlyn Wesley for providing invaluable experimental support for animal studies. We thank Ms. Josie Habak for her long-standing support and advice from an advocate’s perspective, and the International Society of Paediatric Oncology European (SIOPEN) New Drug Development committee for feedback on efficacy testing. Alisertib was kindly supplied by Puma Biotechnologies. The authors thank the Children’s Cancer Institute Animal Facility, the Children’s Cancer Institute Functional High Throughput Technologies and Bioresources and Data Enabling Platforms and NSW Health Statewide Biobank for providing support to this study and acknowledge Compounds Australia and their provision of services for storing and managing the screening libraries. This study was supported by funding from Neuroblastoma Australia (JIF, TNT) and the National Health and Medical Research Council Investigator (APP2017256; MH) and Synergy (APP2018642; MH, MDN) Grants.

