## Supplementary Figures for "Preclinical trial supports dual inhibition of BCL2 and Aurora kinase A for *MYCN*-amplified high-risk neuroblastoma"

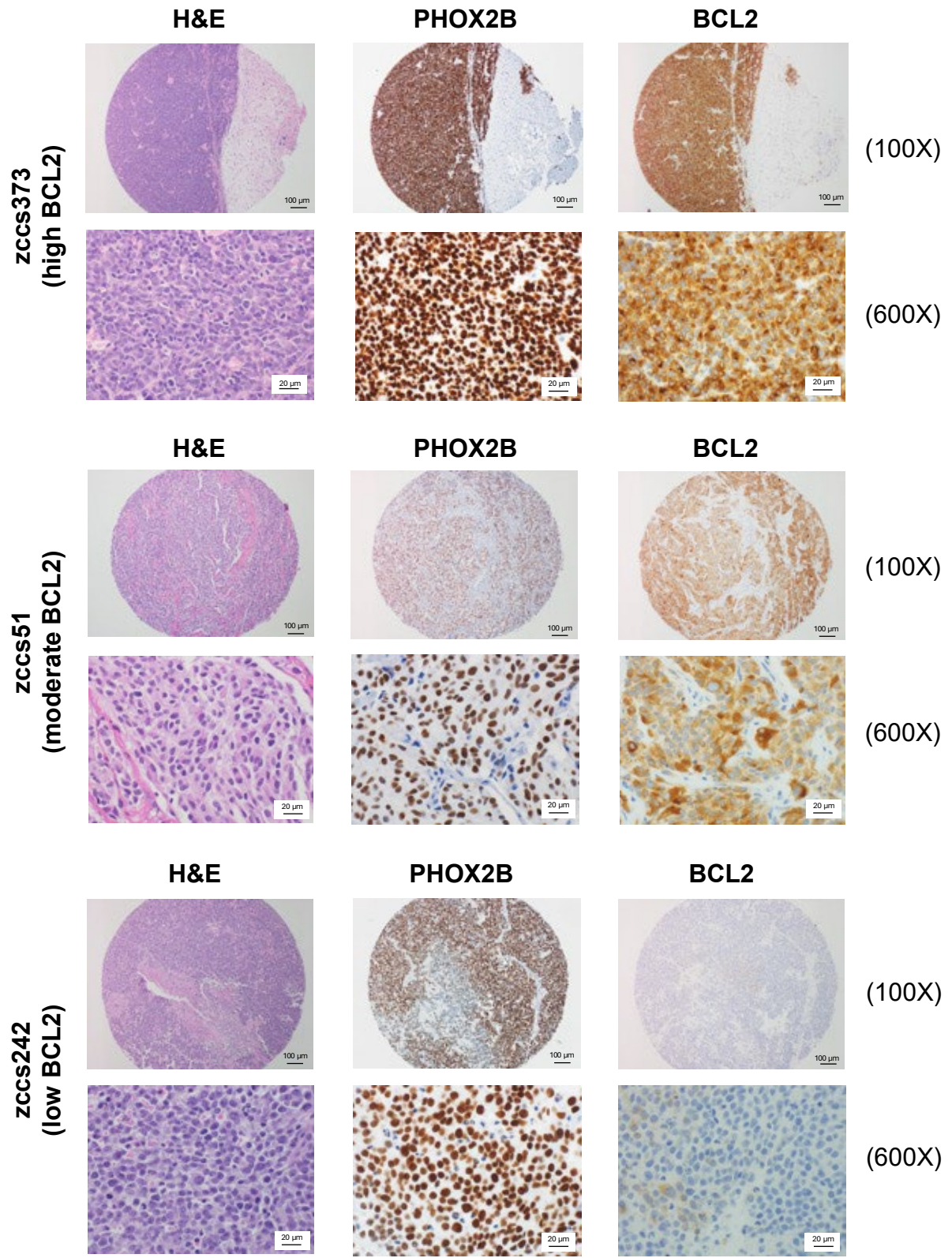

**Supplementary Figure S1.** A tissue microarray (TMA) of neuroblastoma PDX models (n = 34) was stained with H&E and antibodies to PHOX2B (diagnostic marker) and BCL2. Three models with high-, moderate- and low-expression of BCL2 protein are shown. Photos were captured at 100X and 600X magnification. Scale bars represent 100 μm and 20 μm for 100X and 600X magnifications, respectively.

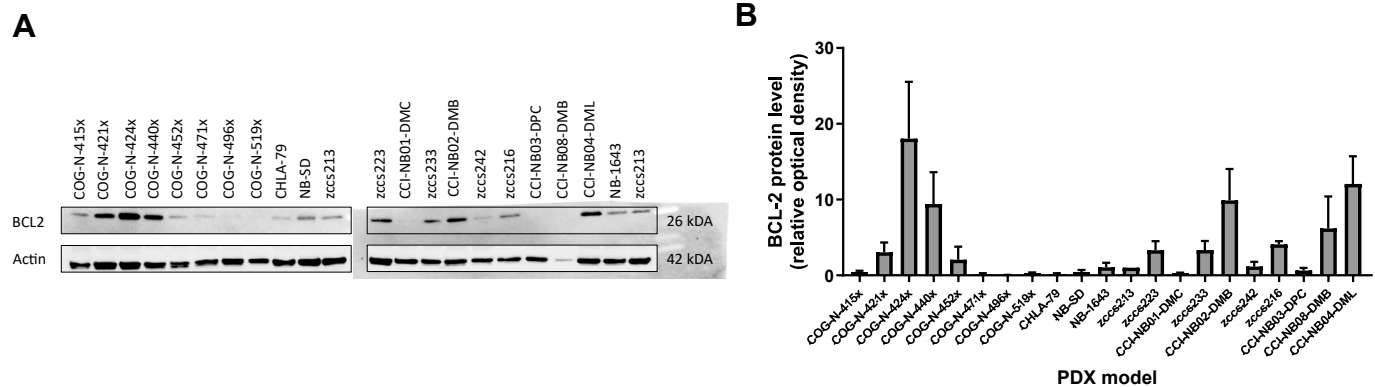

**Supplementary Figure S2. (A)** Western blot of BCL2 protein in selected HR-NB PDX panel (n=21). Beta actin was used as a loading control. **(B)** Relative optical density of BCL2 following normalization to beta actin.

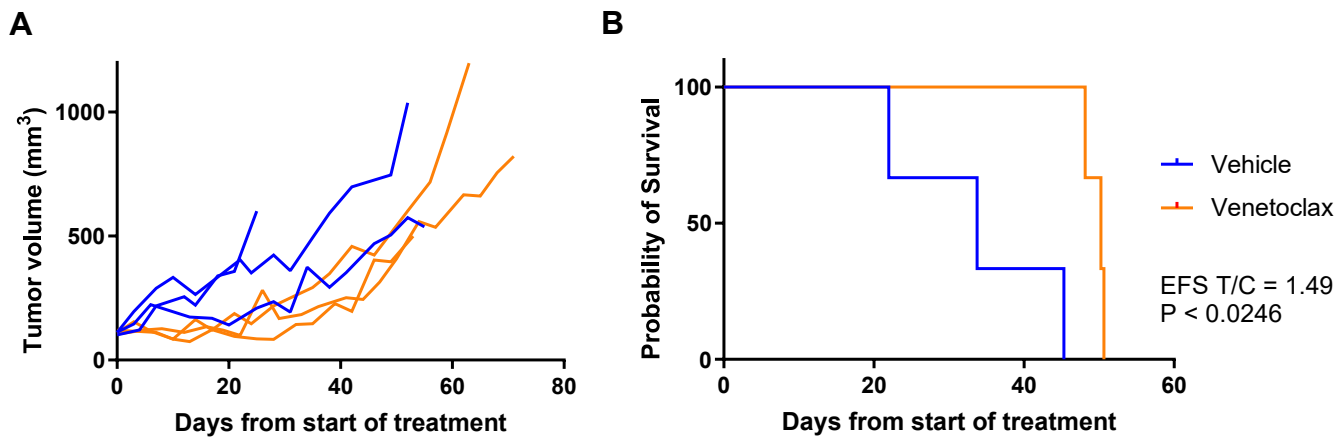

**Supplementary Figure S3.** Growth (A) and survival (B) curves for zccs223 PDX mice following treatment with vehicle or venetoclax monotherapy (n=3 mice per treatment group). EFS T/C and P values are displayed.

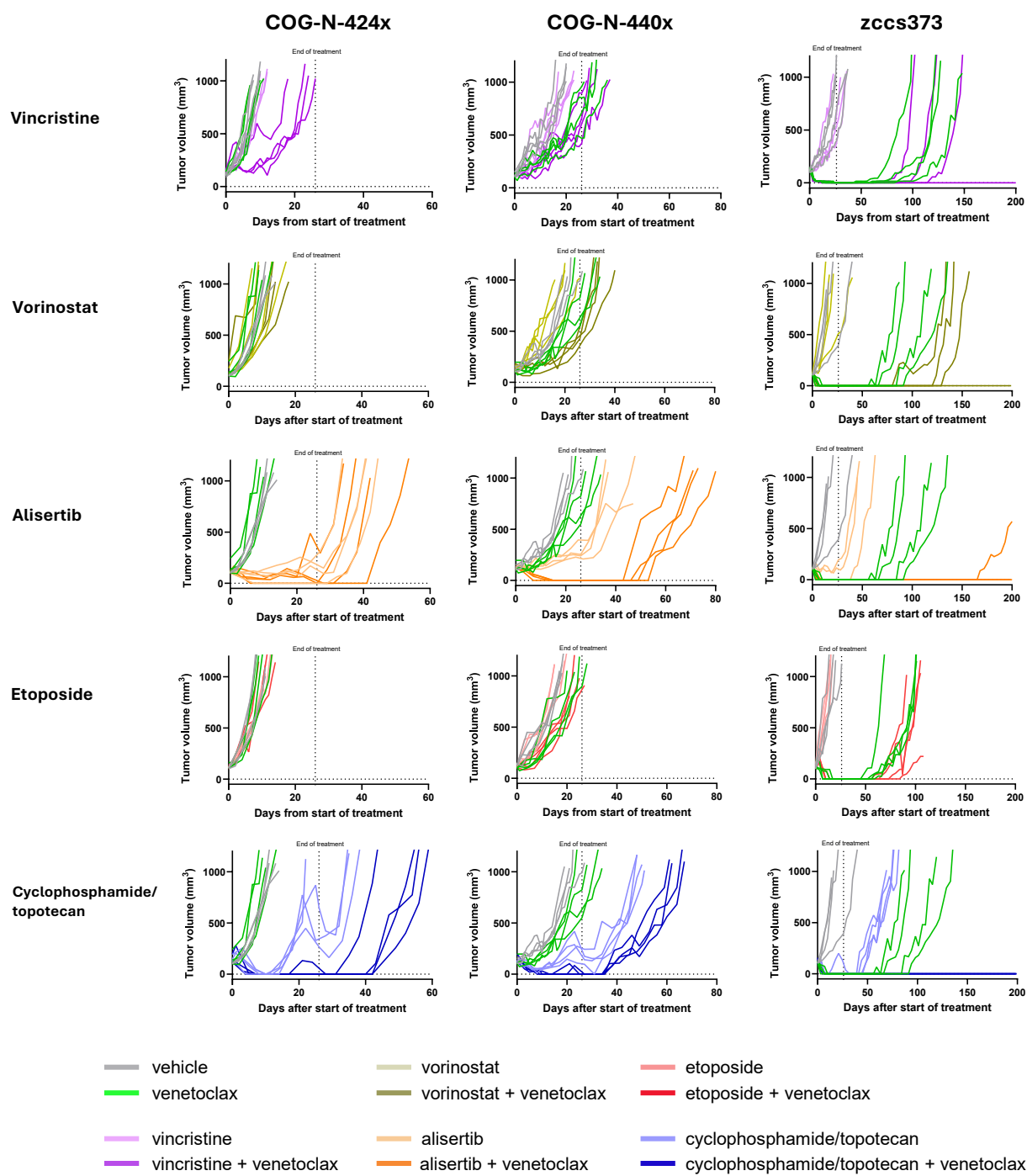

**Supplementary Figure S4.** Growth curve of PDX mice treated with venetoclax combinations.

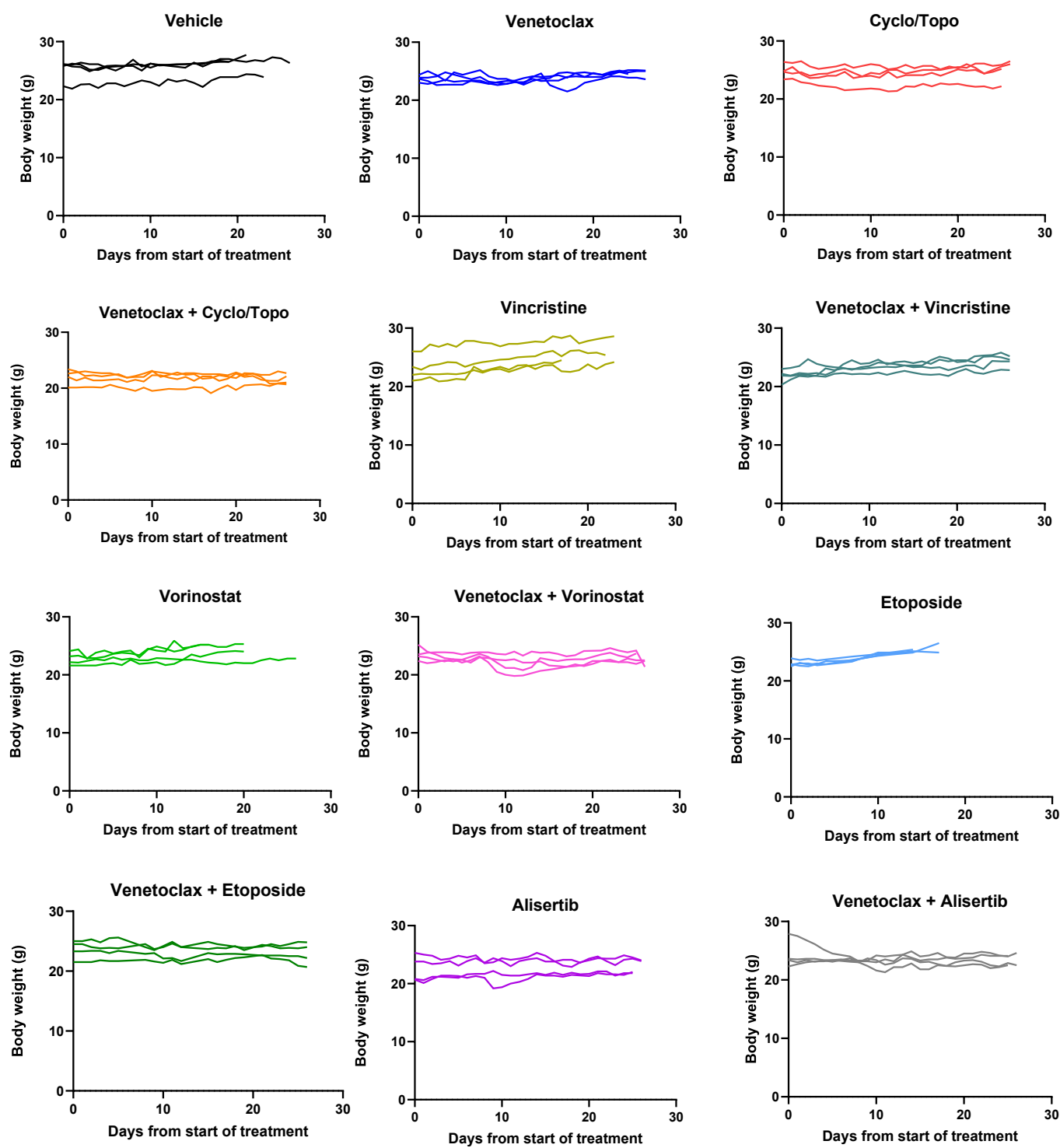

**Supplementary Figure S5.** Mice body weight following treatment with venetoclax combinations and their monotherapies

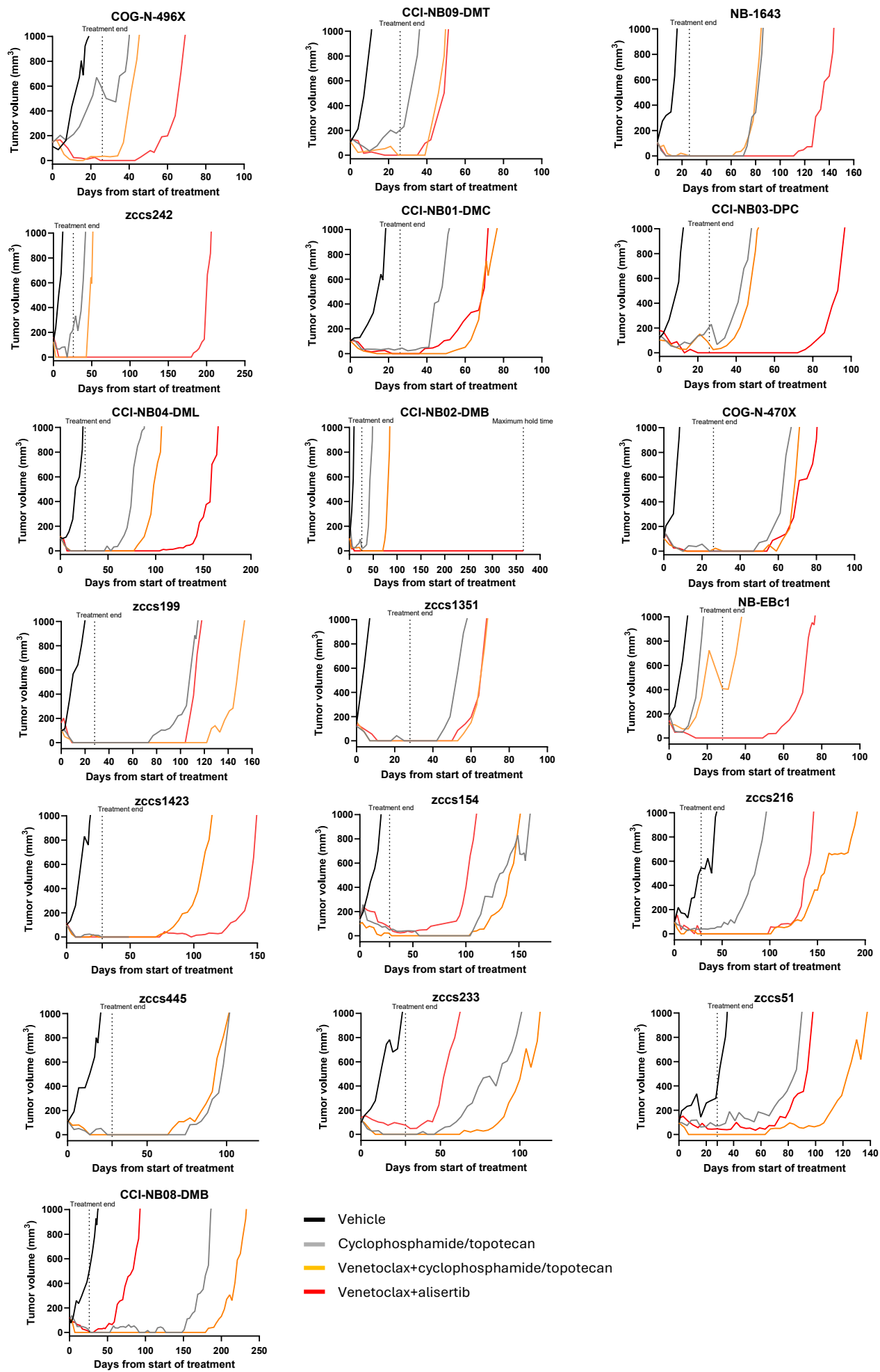

**Supplementary Figure S6.** Growth curve of PDX mice treated with venetoclax combinations in single mouse trial

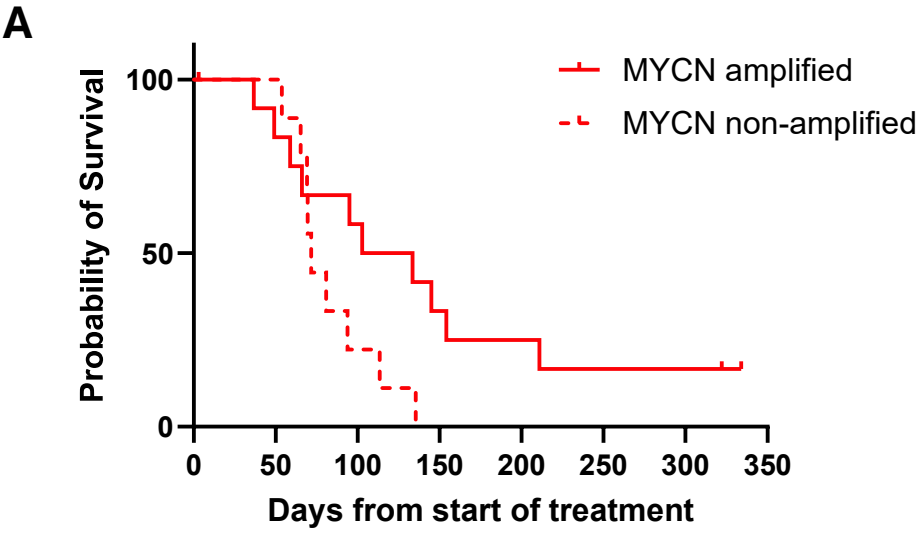

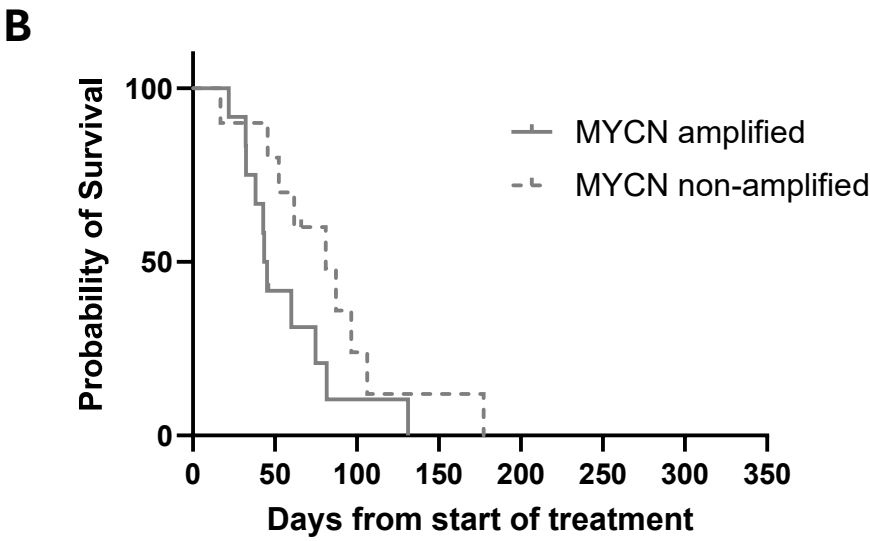

**Supplementary Figure S7.** Survival curve of MYCN amplified and non-amplified PDX models in single mouse trial, treated with **(A)** venetoclax-alisertib or **(B)** standard chemotherapy cyclophosphamide-topotecan.

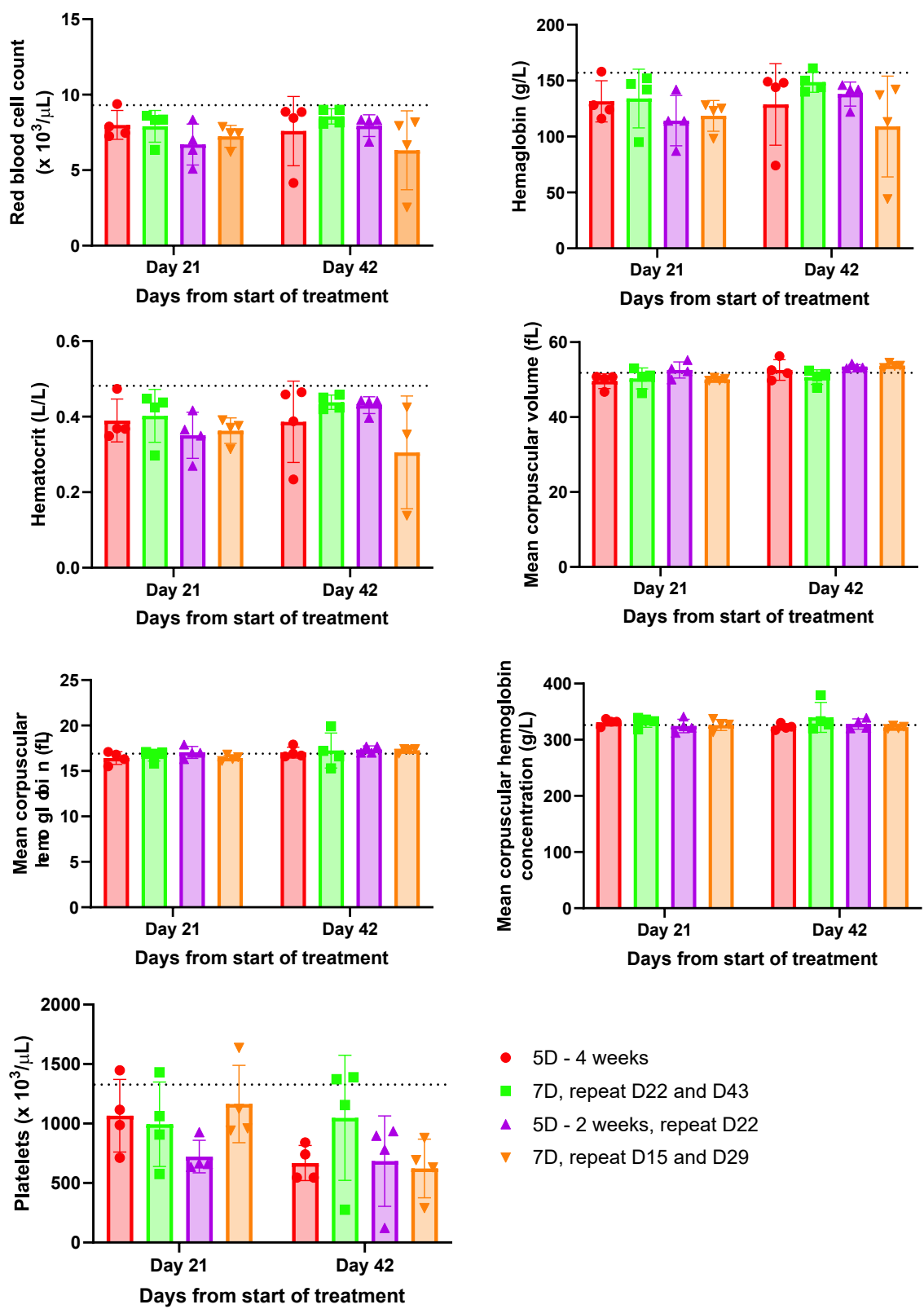

**Supplementary Figure S8.** Hematological parameters in mice treated with different schedules of venetoclax-alisertib combination. Blood was collected at day 21 and day 42 after start of treatment. Dotted line represents the average level of each parameter in untreated NSG mice.

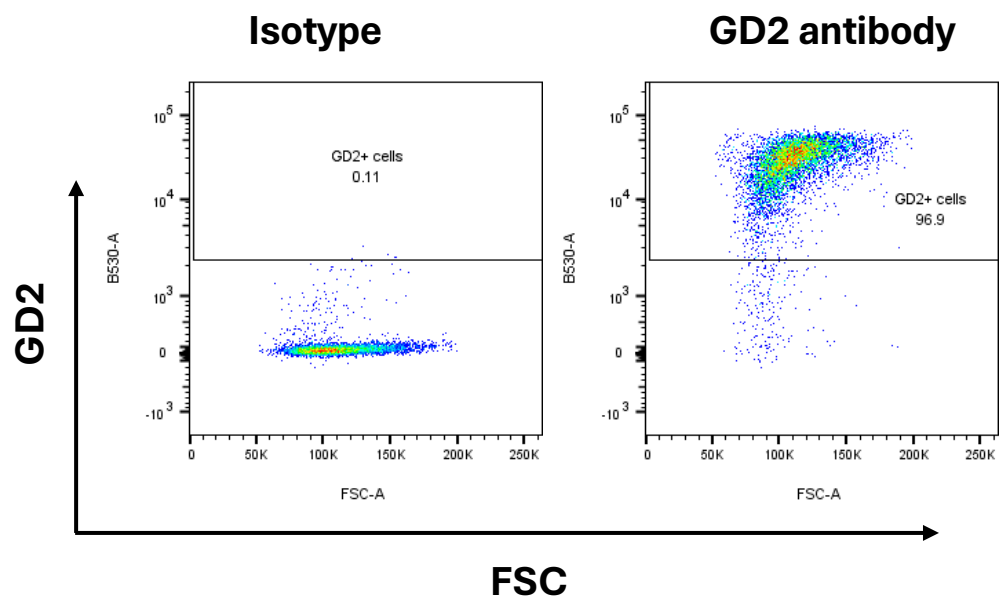

**Supplementary Figure S9.** Level of GD2 expression in COG-N-440x.

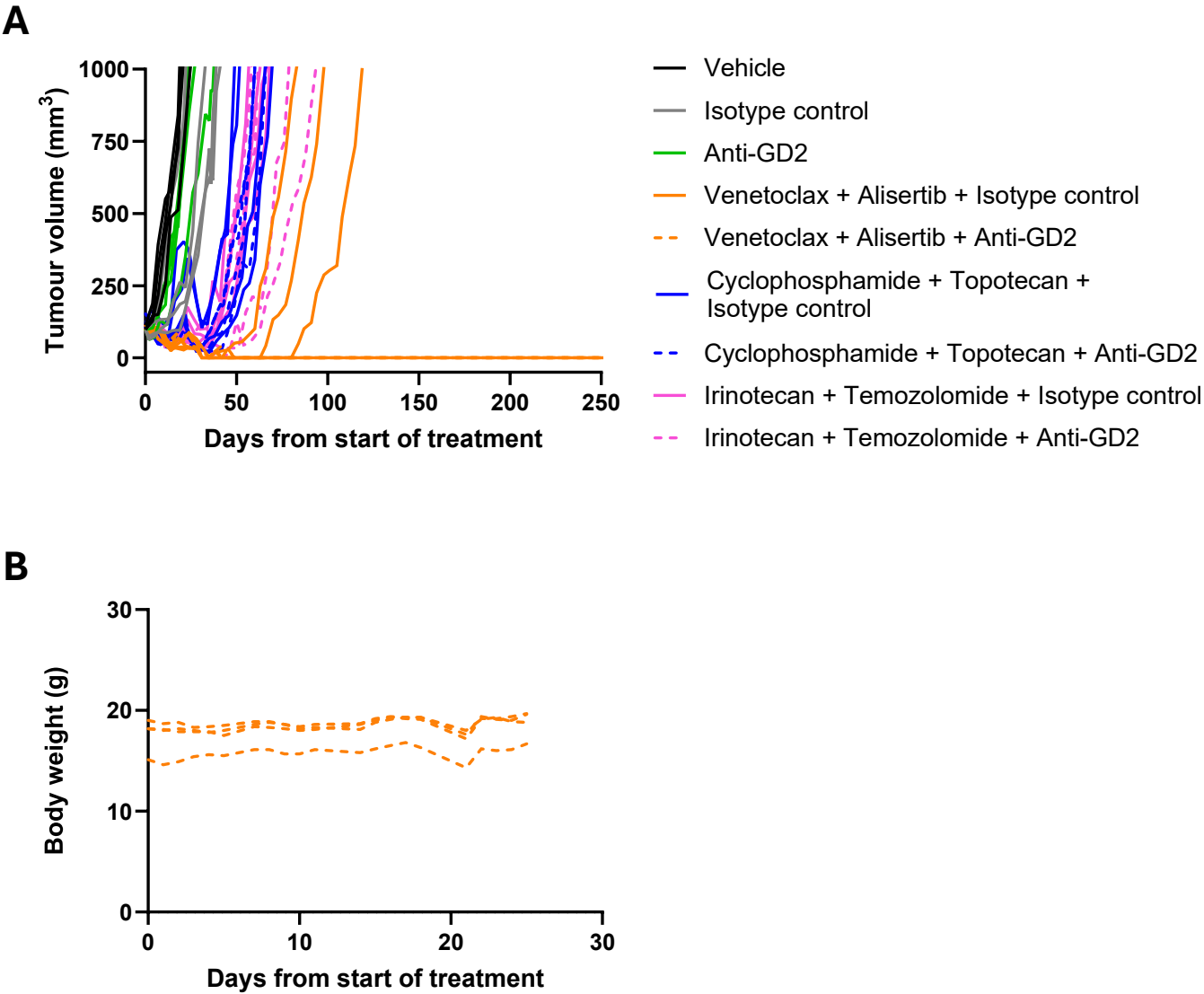

**Supplementary Figure S10. (A)** Tumor growth curve of COG-N-440x model treated with chemo-immunotherapy or venetoclax-alisertib-anti GD2 combination. **(B)** Body weight of mice treated with venetoclax-alisertib-anti-GD2.
